# Novel genetic features of human and mouse Purkinje cell differentiation defined by comparative transcriptomics

**DOI:** 10.1101/2020.01.07.897371

**Authors:** David E. Buchholz, Thomas S. Carroll, Arif Kocabas, Xiaodong Zhu, Hourinaz Behesti, Phyllis L. Faust, Lauren Stalbow, Yin Fang, Mary E. Hatten

**Affiliations:** Laboratory of Developmental Neurobiology, The Rockefeller University, New York, NY 10065, USA; Bioinformatics Resource Center, The Rockefeller University, New York, NY 10065, USA; Department of Pathology and Cell Biology, Columbia University Irving Medical Center and the New York Presbyterian Hospital, New York, NY 10032

**Keywords:** Cerebellum, Purkinje neuron, mitochondria, autophagy, stem cell

## Abstract

Comparative transcriptomics between differentiating human pluripotent stem cells (hPSC) and developing mouse neurons offers a powerful approach to compare genetic and epigenetic pathways in human and mouse neurons. To analyze human Purkinje cell (PC) differentiation, we optimized a protocol to generate hPSC-PCs that formed synapses when cultured with mouse cerebellar glia and granule cells and fired large calcium currents, measured with the genetically encoded calcium indicator jRGECO1a. To directly compare global gene expression of hPSC-PCs with developing mouse PCs, we used translating ribosomal affinity purification (TRAP). As a first step, we used *Tg(Pcp2-L10a-Egfp)* TRAP mice to profile actively transcribed genes in developing postnatal mouse PCs, and used metagene projection to identify the most salient patterns of PC gene expression over time. We then created a transgenic *Pcp2-L10a-Egfp* TRAP hESC line to profile gene expression in differentiating hPSC-PCs, finding that the key gene expression pathways of differentiated hPSC-PCs most closely matched those of late juvenile, mouse PCs (P21). Comparative bioinformatics identified classical PC gene signatures as well as novel mitochondrial and autophagy gene pathways during the differentiation of both mouse and human PCs. In addition, we identified genes expressed in hPSC-PCs but not mouse PCs and confirmed protein expression of a novel human PC gene, CD40LG, expressed in both hPSC-PCs and native human cerebellar tissue. This study therefore provides the first direct comparison of hPSC-PC and mouse PC gene expression and a robust method for generating differentiated hPSC-PCs with human-specific gene expression for modeling developmental and degenerative cerebellar disorders.

**Significance Statement:** To compare global gene expression features of differentiating human pluripotent stem cell-derived Purkinje cells (hPSC-PC) and developing mouse Purkinje cells (PC) we derived hPSC-PCs and compared gene expression datasets from human and mouse PCs. We optimized a differentiation protocol that generated hPSC-PCs most similar in gene expression to mouse P21 PCs. Metagene projection analysis of mouse PC gene expression over postnatal development identified both classical PC marker genes as well as novel mitochondrial and autophagy gene pathways. These key gene expression patterns were conserved in differentiating hPSC-PCs. We further identified differences in timing and expression of key gene sets between mouse and hPSC-PCs and confirmed expression of a novel human PC marker, CD40LG, in human cerebellar tissue.

## Introduction

Emerging evidence supports a role for the cerebellum in wide range of cognitive functions, including language, visuospatial memory, attention, and emotion, in addition to classical functions in adaptive, feed forward motor control (1–3). Cerebellar defects therefore contribute to a broad spectrum of neurological disorders including ataxias, autism spectrum disorder (ASD), intellectual disability, and other cerebellar-based behavioral syndromes (2–7). As the sole output neuron of the cerebellar cortex, the Purkinje cell (PC) plays a key role in both development and function of the cerebellum, integrating information from their primary inputs, the cerebellar granule cells (GCs) and climbing fiber afferents (8). A loss of PCs is one of the most consistent findings in postmortem studies in patients with ASD (4), and specific targeting of PCs in mouse models of ASD-associated genes leads to impaired cerebellar learning (9) and social behaviors (6, 10). Notably, PC degeneration is also a hallmark of human spinocerebellar ataxias (7).

While modeling genetic disorders in mice has provided fundamental insights into disease mechanisms, human disease often cannot be fully recapitulated in the mouse. A prominent example is ataxia-telangiectasia, which shows massive loss of PCs in humans but not in the mouse (11). Human pluripotent stem cells (hPSCs) provide a complementary approach to studying human disease in the mouse (12–14). Validated methods to derive specific neural subtypes from hPSCs are necessary prerequisites to disease modeling. We and others have recently developed protocols to derive PCs from hPSCs (15–18).

One limitation of most hPSC-derived CNS neurons is the lack of genetic information, especially of transcriptomic signatures, to rigorously identify specific types of neurons and to compare their development across species. Here, we present an optimized method to generate well-differentiated hPSC-PCs, and use translating ribosomal affinity purification (TRAP) to directly compare global gene expression patterns of developing mouse PCs with that of differentiating hPSC-derived PCs (hPSC-PCs). After induction with signals that induce PCs, purification and co-culture with cerebellar glia, hPSC-PCs formed synapses with mouse granule cells and fired large calcium currents, measured with the genetically encoded calcium indicator jRGECO1a. Metagene projection analysis of global gene expression patterns revealed that differentiating hPSC-PCs share classical and novel developmental gene expression signatures with developing mouse PCs that include mitochondrial and autophagy gene pathways. Comparative bioinformatics of key gene pathways showed that hPSC-PCs closely match juvenile P21 mouse PCs, suggesting that they are relatively mature. Gene expression profiling also identified human-specific genes in hPSC-PCs. Protein expression for one of these human specific genes, CD40LG, a TNF superfamily member, was confirmed in both hPSC-PCs and native human cerebellar tissue. This study therefore provides the first direct comparison between hPSC-PC and mouse PC gene expression.

## Results

### Differentiation of hPSCs to PCs

We previously reported a protocol to generate cerebellar PCs from hPSCs, which employed an approach that has proven broadly successful for hPSC differentiation: recapitulation of development through the addition of inductive signals (18, 19). Here we have optimized that protocol, quantifying the effect of signaling molecules on early differentiation and introducing new methods for isolation of immature PCs and co-culture with mouse cerebellar glia and granule cells. These additions provide a robust method for generation of hPSC-PCs.

Neural induction was achieved by culture of hPSC aggregates in NOGGIN (20) and nicotinamide (21, 22). Nicotinamide significantly decreased expression of the pluripotency marker *OCT4* while significantly increasing expression of the mid/hindbrain neural tube markers *EN1* and *EN2* after 6 days of differentiation when combined with additional differentiation factors (Figures 1B and S1A/B). Specific concentrations of GSK3β inhibitors such as CHIR99021 have previously been used to mimic developmental Wnt signaling gradients to direct hPSCs to specific rostral-caudal domains, giving rise to midbrain dopaminergic neurons (23–25). We confirmed that increasing concentrations of CHIR99021 led to progressive caudalization of neural tissue in our differentiation protocol, with a concentration of 1.5μM generating the highest expression of cerebellar anlage markers *EN1, EN2*, and *GBX2* after 6 days when combined with additional differentiation factors (Figures 1B and S1A/B). Following neural induction and initial rostral-caudal patterning, the cerebellum is subsequently specified by high levels of FGF8b signaling (8). We found that addition of 100ng/mL FGF8b at day 4 of differentiation led to a significant increase in *En1, En2*, and *Gbx2* expression at day 6 compared to 1ng/mL FGF8b (Figures 1B and S1A). The addition of NOGGIN, nicotinamide, CHIR99021, and FGF8b at specific concentrations and times led to nearly uniform induction of midbrain/hindbrain character by day 6, defined by co-expression of *EN1/OTX2* (Figure 1B). After plating the hPSC-aggregates on laminin at day 6 and continued culture in FGF8b, our cultures separated into *EN1*^+^/*OTX2*^+^ (midbrain) and *EN1*^+^/*OTX2*^−^ (hindbrain/cerebellum) regions (Figure 1C). FGF8b was removed on day 12 and BDNF was added to support post-mitotic neurons. From day 12-24 neural rosettes expressing the cerebellar ventricular zone marker KIRREL2 gave rise to increasing numbers of adjacent cells outside of the rosettes expressing the earliest post-mitotic PC marker, CORL2/SKOR2 (Figure 1D), similar to organization within the developing cerebellum (8). The definitive post-mitotic Purkinje cell marker PCP2 was observed starting at day 18 onwards (Figure 1E).

**Figure 1.**
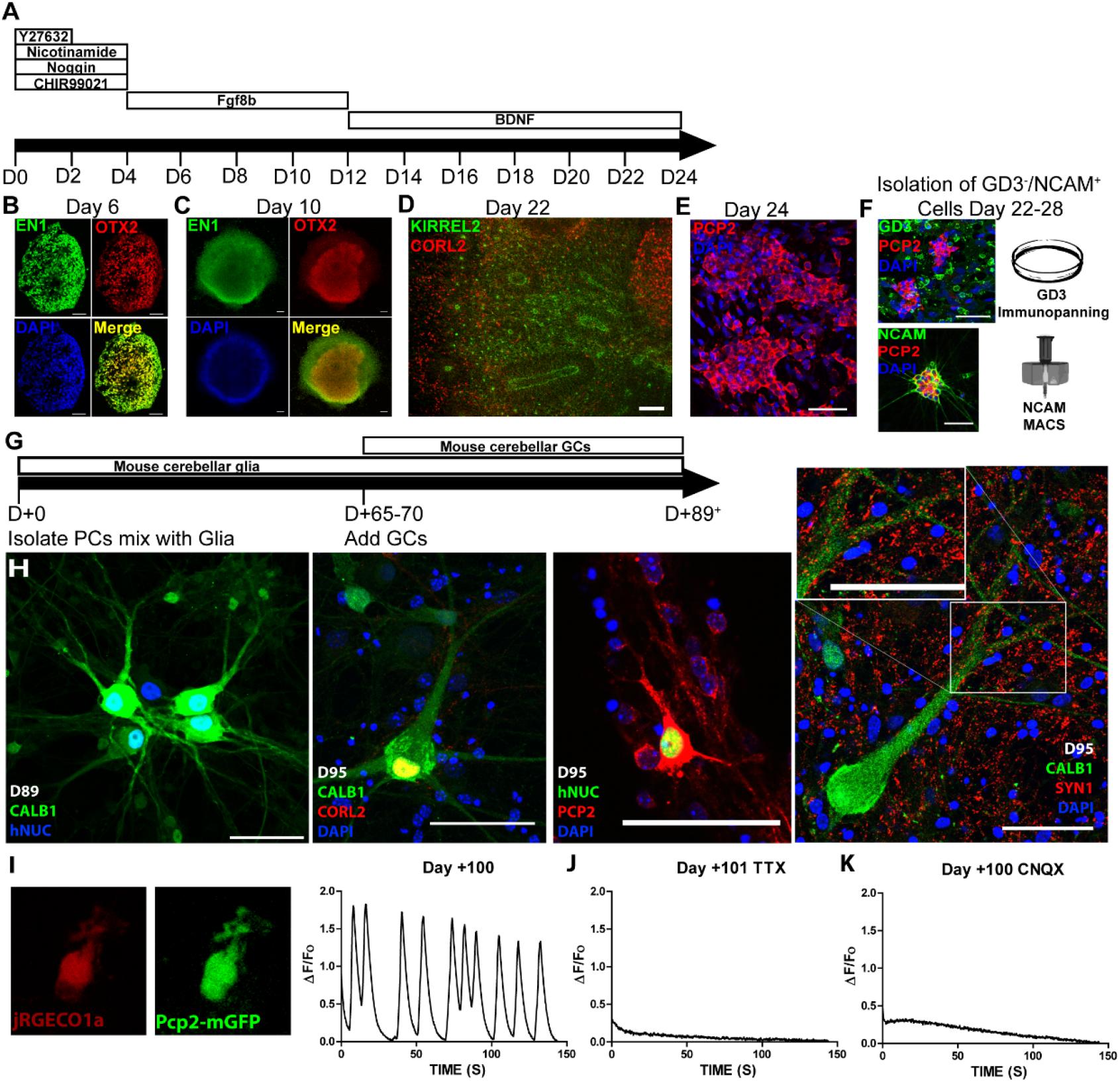
Differentiation of hPSCs to PCs. **A.** Schematic of the first phase of differentiation. **B.** Immunolabeling of a cryo-section of a neural aggregate after 6 days of differentiation. 10 μm z-projection. Scale bar 50μm. **C.** Immunolabeling of an attached neural aggregate after 10 days of differentiation. Scale bar 100μm. **D.** Immunolabeling of an attached neural aggregate after 22 days of differentiation. 18 μm z-projection. Scale bar 100 μm. **E.** Immunolabeling of an attached neural aggregate after 24 days of differentiation. Scale bar 50 μm. **F.** Immunolabeling of cell-surface markers used for isolation of hPSC-PCs from differentiated cultures. Scale bars 50 μm. **G.** Schematic of the second phase of differentiation. **H.** Immunolabeling of hPSC-PCs after isolation and co-culture with mouse glia and granule cells for an additional >89 days. From left to right: 7 μm z-projection, 5 μm z-projection, 5 μm z-projection, single optical section. Scale bars 50μm. hNUC – human nuclear antigen. **I.** Live imaging of genetically encoded calcium indicator jRGECO1a and hPSC-PC reporter Pcp2-mGFP after 100 days of co-culture with mouse glia and granule cells and a trace of the change in jRGECO1a fluorescence (ΔF/Fo) over time. **J.** Representative trace of jRGECO1a fluorescence in the presence of the Na^+^ channel antagonist tetrodotoxin (TTX) after 101 days of co-culture with mouse glia and granule cells. **K.** Representative trace of jRGECO1a fluorescence in the presence of the glutamate receptor antagonist CNQX after 100 days of co-culture with mouse glia and granule cells.

To provide a more defined model system, we developed a two-step procedure to purify PCs after 22-28 days of differentiation: negative selection by GD3 immunopanning (26) and positive selection by NCAM1 magnetic cell sorting (Figure 1F). Since the maturation of mouse PCs is strongly dependent on cell-cell interactions with glia and cerebellar GCs (26), we first co-cultured hPSC-PCs with cerebellar glia for 65-70 days followed by co-culture with their presynaptic partner neurons, GCs, for 25-30 days (Figure 1G). Culturing purified hPSC-PCs with glia prior to culture with GCs resulted in more robust expression of CALB1 than culturing with culturing the cells with GCs alone (Figure S1C). In addition, after co-culture with glia, hPSC-PCs co-cultured with mouse GCs extended thick primary dendrites similar to those seen in developing human PCs (Figure S4B; Rakic and Sidman, 1970; Zecevic and Rakic, 1976) and expressed other classical mouse PC marker genes, including CORL2/SKOR2, and PCP2 (Figure 1H). Immunolabeling of the presynaptic protein SYN1 revealed numerous puncta along hPSC-PC dendrites, indicative of the formation of synapses between hPSC-PCs and GCs (Figure 1H) (27, 28). After ∼17 weeks of differentiation, the morphology of hPSC-PCs resembled that of immature human fetal PCs after ∼18-25 weeks of gestation (Figure S4B).

### Functional testing of hPSC-PCs with genetically encoded calcium indicators

To confirm the presence of functional synaptic connections between mouse GCs and hPSC-PCs, we nucleofected the genetically encoded calcium indicator jRGECO1a (29) into freshly purified hPSC-PCs (Figure 1I). After co-culture with mouse cerebellar glia for 70 days and GCs for an additional 30 days, several different calcium firing patterns were observed in labeled Pcp2^+^ hPSC-PCs, (Figures 1I and S1D, and movies S1-3). A slow decay of 30-90 seconds was observed in many responses, suggesting release of calcium from intracellular stores (Figure S1D). Signals greater than 1-fold change in fluorescence over baseline (ΔF/F_0_) were only observed in the presence of granule cells, when SYN1^+^ synapses had been observed between GCs and hPSC-PCs. hPSC-PC activity was inhibited by addition of the voltage-gated sodium channel antagonist tetrodotoxin and by addition of the ionotropic glutamate receptor antagonist CNQX (Figure 1J and 1K). Inhibition by CNQX is consistent with the formation of functional synapses between glutamatergic GCs and hPSC-PCs.

### Determination of hPSC-PC maturity by comparative transcriptomics

To examine the identity of hPSC-PCs at the transcriptome level and to compare global patterns of gene expression of differentiating hPSC-PCs with developing mouse PCs, we used TRAP to purify mRNAs (30). Lentiviral delivery of the L10a-EGFP tag under control of the *Pcp2* promoter (31) generated a Pcp2-L10a-Egfp hPSC line (Figures 2A and S2A/B). By immunofluorescence, L10a-EGFP expression was restricted to PCP2^+^ hPSC cells that co-expressed the classical PC marker CALB1 (Figure 2A). L10a-EGFP properly localized to the soma and nucleolus, providing a convenient live tag for differentiating hPSC-PCs (Figures 2A and S2B) (32). Some PCP2^+^ cells did not express the L10-EGFP tag, suggesting partial silencing of the lentiviral construct, in agreement with prior reports of lentiviral transgenesis in hPSCs (33). Translating mRNA was affinity purified from hPSC-PCs after 24 days of differentiation, and after co-culture with mouse cerebellar glia and GCs for an additional +95 days and analyzed by RNAseq (Figure 2B). Principal component analysis of RNAseq datasets from undifferentiated hPSC lines H1 and RUES2 and hPSC-PCs on day 24 or day +95 showed tight clustering within replicates and separation between groups (Figure 2C).

**Figure 2.**
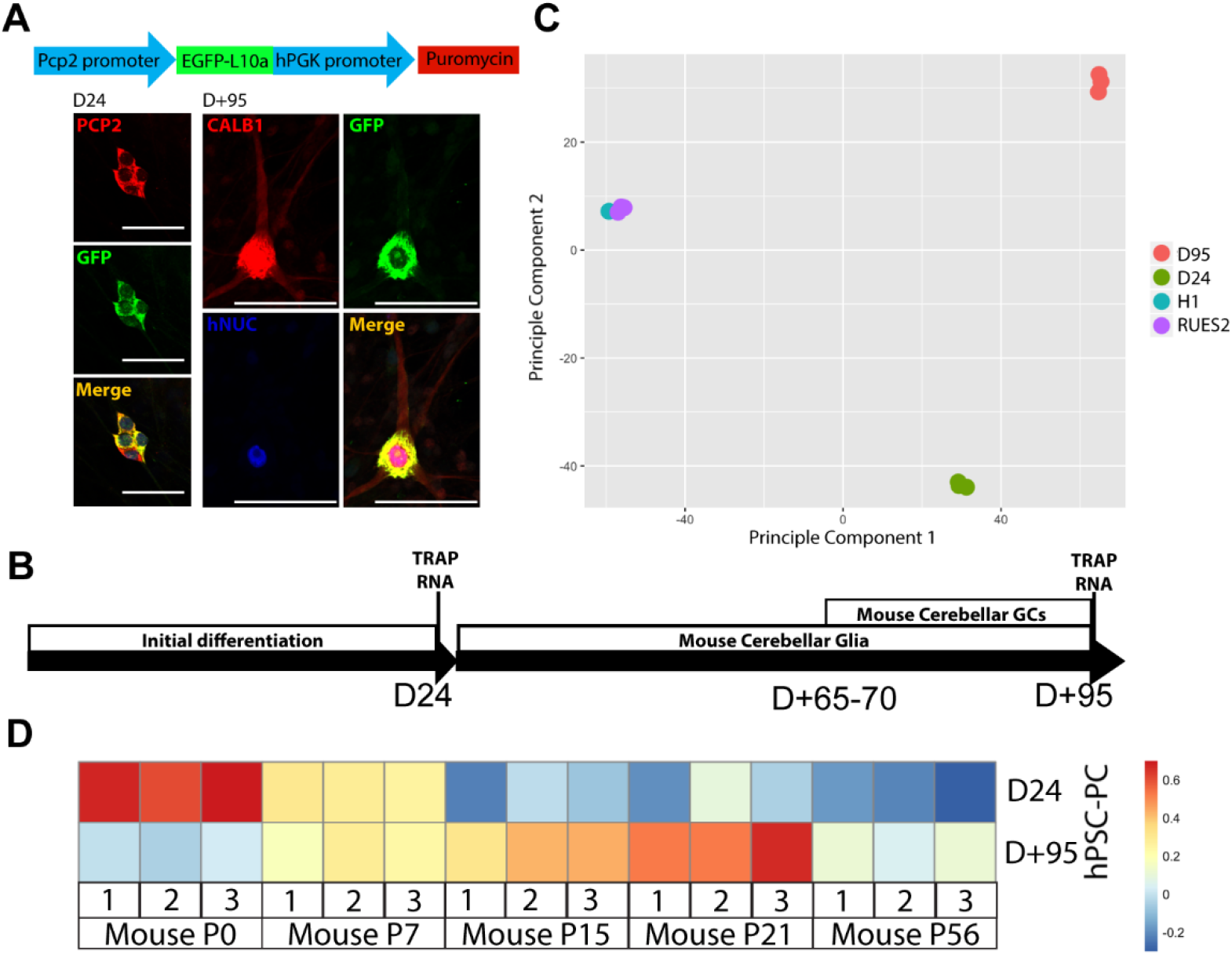
Comparison of hPSC-PC gene expression to mouse PC gene expression over development. **A.** Schematic of the lentiviral construct used to create the PCP2-EGFP-L10a TRAP hPSC line and immunolabeling of the construct in differentiating hPSC-PCs. 4 μm z-projections. Scale bars 50μm. **B.** Schematic depicting timing of TRAP RNA isolation from differentiation hPSC-PCs. **C.** Principle component analysis of differentiating hPSC-PCs after 24 days and +95 days as well as undifferentiated hESC lines (H1, RUES2). **D.** Heat map depicting expression levels in mouse PCs over postnatal development of gene sets defined as log2 four-fold change between Day 24 and Day +95 in hPSC-PCs. Day 24 hPSC-PCs are most similar to P0 mouse PCs (p=0.0014). Day +95 PCs are most similar to P21 mouse PCs (p=1.74^−32^).

To profile global gene expression patterns of mouse PCs at different stages of development, we used the *Tg(Pcp2-L10a-Egfp)* TRAP mouse line (Figure S2C). mRNAs were purified at P0, when mouse PCs are beginning to differentiate; at P7, when GCs are migrating and extending parallel fibers that synapse with nascent PCs; at P15, when PCs are maturing; at P21, when the cerebellar circuitry has formed and PC dendritic arbors are nearly mature; and at P56, when PCs are fully mature (Figure S4A). Isolated mRNA was analyzed by microarray. RNAseq gene sets defined in hPSC-PCs were used to compare gene expression between species and platforms. Gene sets for hPSC-PC time points were defined as log2 four-fold difference between day 24 and day +95. Expression levels of hPSC-PC day 24 and day +95 gene sets were analyzed in mouse PC expression data (Figure 2D). hPSC-PCs at day 24 were most similar to mouse PCs at P0 (p=5.26×10^−17^), while hPSC-PCs at day +95 were most similar to mouse PCs at P21 (p=3.73×10^−5^). For added stringency, we processed the data through an *in silico* combined human-mouse reference genome to subtract possible mouse reads and observed the same patterns (Figure S2D) (34). Defining gene sets in mouse PCs (100 highest expressed genes per time point) and assessing expression levels in hPSC-PCs also gave the same pattern (Figures S2E and S2F). These data indicate that at the transcriptome level, hPSC-PCs mature over differentiation to a stage most similar to juvenile mouse P21 PCs.

### Non-negative matrix factorization defines developmental PC gene signatures

To assess global transcription profiles of mouse PCs over postnatal developmental and identify key gene pathways, we used a non-negative matrix factorization (NMF) approach to define clusters of genes with similar developmental expression patterns, termed metagenes (35–37). This method revealed several metagenes with high cophenetic correlation coefficients (Figure S3A). Of these, number five (five metagenes) had the lowest dispersion and was therefore used in this study (Figures 3A and S3A). We then identified gene ontology (GO) sets with a correlation coefficient >0.85 to any metagene (Figure 3B). GO terms associated with PC development were highly correlated with metagene 1 (Figures 3B and S3B), validating our NMF approach. The level of expression of PC developmental genes in this metagene set increased in mouse PCs from P0 to P7 and also in hPSC-PCs from day 24 to day +95. Thus, at the transcriptome level, hPSC-PCs expressed gene sets that identify developing mouse PCs.

**Figure 3.**
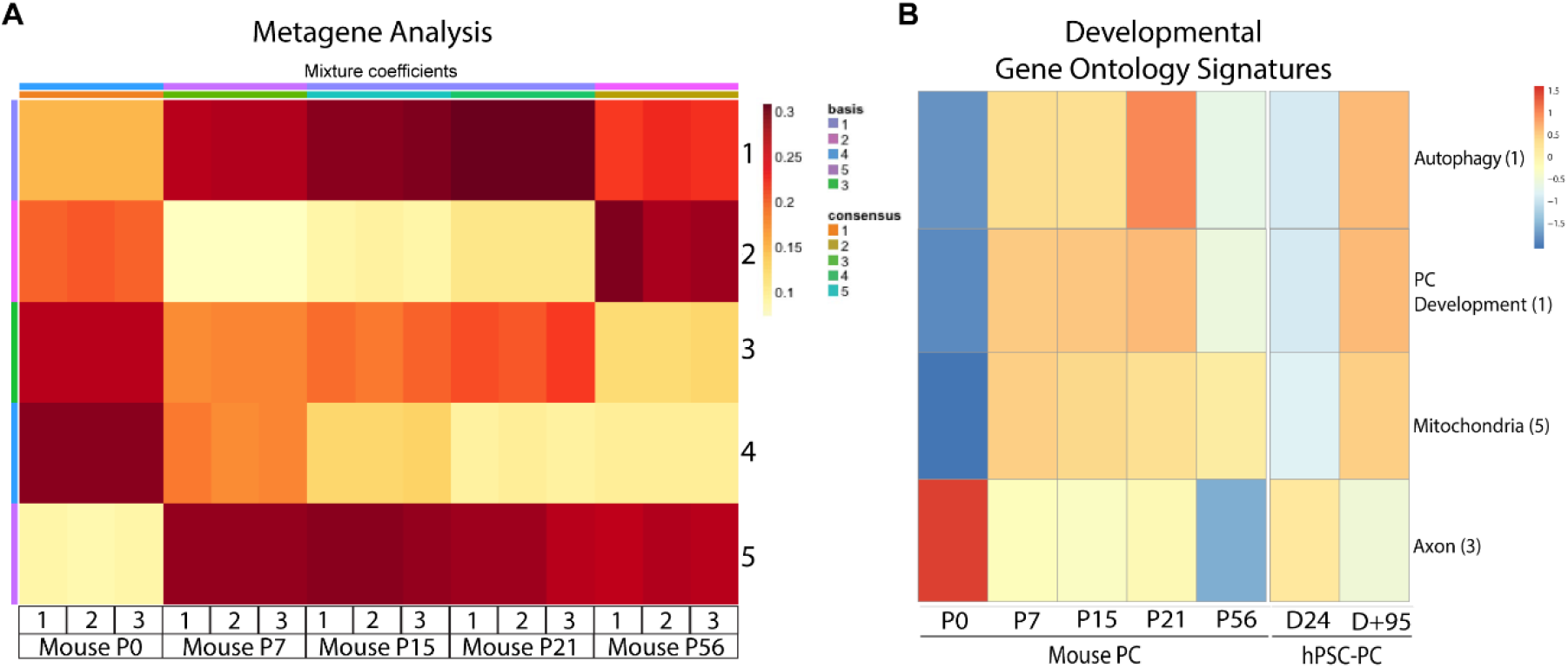
NMF analysis of mouse PC developmental gene expression and comparison to hPSC-PCs. **A.** Metagenes defined by NMF analysis of mouse PC gene expression over postnatal development. **B.** Gene ontology signatures with a correlation coefficient to a metagene in A. of >0.85. Listed next to the terms is the metagene with highest correlation.

At P0, mouse PCs highly expressed gene sets associated with GO terms for axon outgrowth. These gene sets correlated with metagene 3, showing high expression at P0 but then dropping off at P7 and further dropping at P56 (Figures 3B and S3B). The patterns for axon outgrowth similarly decreased in hPSC-PCs from day 24 to day +95. Developmental gene signatures for mitochondria and autophagy correlated highly with metagenes 1 and 5, showing a sharp increase in expression from P0 to P7 (Figures 3B and S3B). Both signatures similarly showed increases in expression in hPSC-PCs from day 24 to day +95. In summary, we observed P0-P7 to be a highly dynamic period for PC gene expression, showing downregulation of axon outgrowth genes and upregulation of mitochondria, autophagy, and classical PC marker genes. These gene expression dynamics were conserved in hPSC-PCs from day 24 to day +95.

### Differences between hPSC-PCs and mouse PCs

Three types of differences in gene expression patterns were observed between mouse PCs and hPSC-PCs. First, the timing of expression of some gene pathways was delayed in hPSC-PCs. Notably, the timing of expression of the thyroid hormone signaling pathway, a key molecular pathway that drives mouse PC dendritic maturation (38) was delayed relative to mouse PCs (Figure 4A). This included delayed expression of *THRA, PPARGC1A*, and *RORα*. This observation is consistent with the slow time course of morphological maturation of hPSC-PCs relative to mouse PCs.

**Figure 4.**
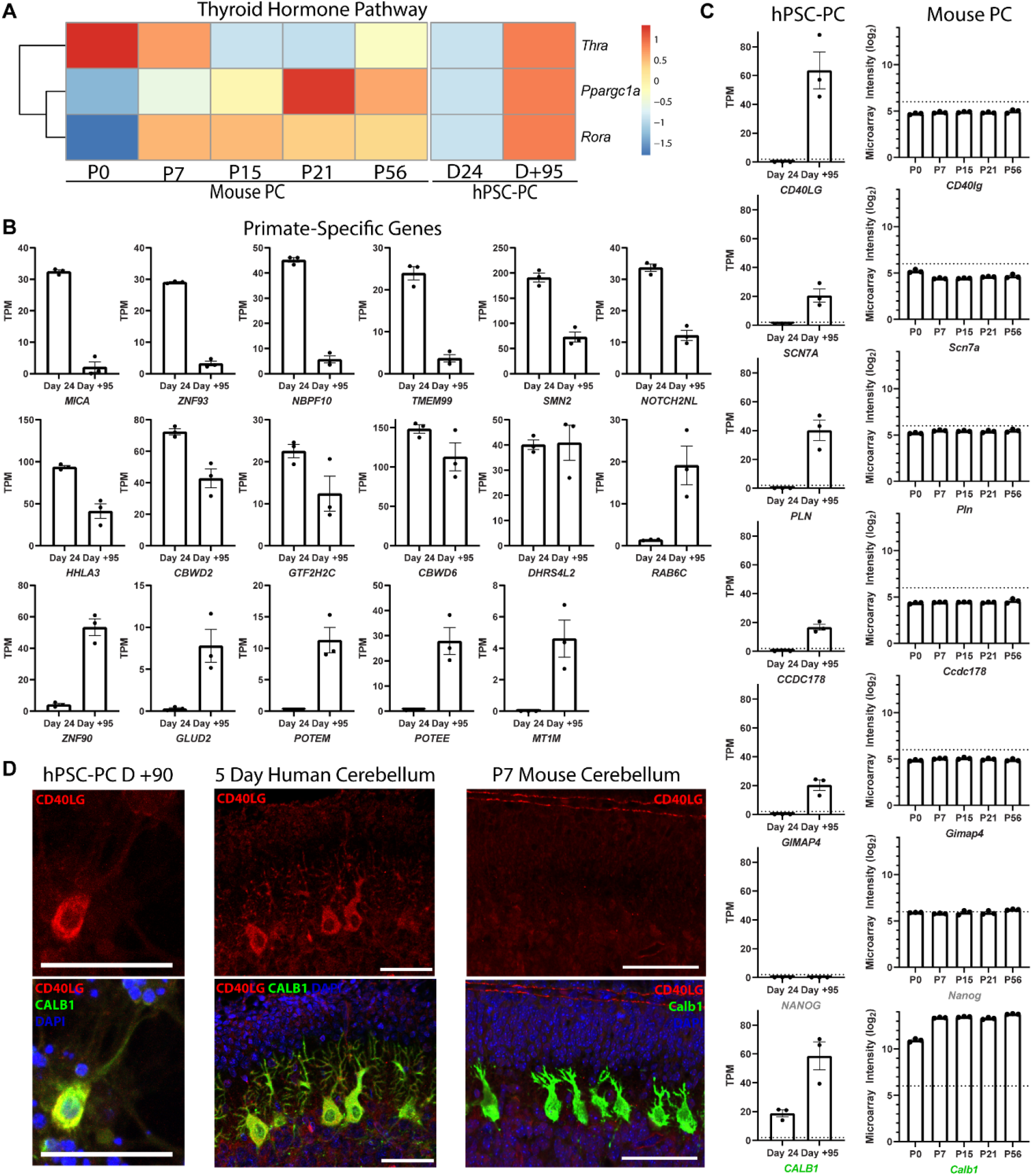
Differences between hPSC-PCs and mouse PCs. **A.** Expression patterns of thyroid hormone signaling pathway. **B.** Expression levels of primate-specific genes in hPSC-PCs at Day 24 and Day +95. Error bars are standard error of the mean **C.** Expression levels of genes upregulated in hPSC-PCs but not expressed in mouse PCs. For hPSC-PCs, background was set at 2 transcripts per million (TPM, dotted line). For mouse PCs, background was set at microarray intensity of 6 (dotted line). The negative control NANOG and the positive control CALB1 are shown for reference. Error bars are standard error of the mean. **D.** Expression of CD40lg in hPSC-PCs, 5 day human cerebellum, but not P7 mouse cerebellum. Scale bars 50μm.

Second, we observed expression of a number of primate-specific genes that do not exist in the mouse genome (Figure 4B). Primate specific genes marked both early and late stages of hPSC-PC differentiation. *MICA, ZNF93, NBPF10, TMEM99, SMN2, NOTCH2NL, HHLA3*, and *CBWD2* were expressed at higher levels at day 24 than at day +95. *GTF2H2C, CBWD6*, and *DHRS4L2* were expressed at relatively similar levels at each time point. *RAB6C, ZNF90, GLUD2, POTEM, POTEE*, and *MT1M* were all upregulated over differentiation from day 24 to day +95. While the roles of many of these genes unclear, *NOTCH2NL* and *GLUD2* have been linked to human-specific changes in brain development (39–45).

Third, we identified genes that are highly upregulated in hPSC-PCs over differentiation but are not detected in postnatal mouse PCs (Figure 4C). These included *CD40LG, SCN7A, PLN, CCDC178*, and *GIMAP4*. We further investigated expression of *CD40LG*, for which antibodies exist to confirm protein expression by immunolabeling. Immunolabeling of hPSC-PCs at day +90 and mouse PCs at P7 confirmed our transcriptomic data, showing co-localization of CD40LG with CALB1 in hPSC-PCs but not mouse PCs (Figure 4D). To verify that CD40LG is expressed in human PCs and not just hPSC-PCs, we labeled newborn human cerebellar tissue. Positive immunolabeling in CALB1^+^ human PCs confirmed that our hPSC-PC model system had captured a species difference in gene expression and identified a novel human-specific PC marker (Figures 4D and S4C).

## Discussion

To provide a more robust model for human Purkinje cell (PC) differentiation, we optimized a protocol to generate hPSC-PCs. A key element of this protocol was the use of MACS to purify immature hPSC-PCs, followed by co-culturing the cells with mouse cerebellar glia and then mouse GCs. With this methodology, hPSC-PCs had a more differentiated morphology, formed SYN1+ synapses with mouse granule cells, and fired large calcium currents, measured with the genetically encoded calcium indicator jRGECO1a. To directly compare global gene expression of hPSC-PCs with developing mouse PCs, we used TRAP, expressing the Pcp2-L10a-Egfp TRAP tag in hPSCs. Comparison of key gene pathways between developing mouse PCs and differentiating hPSC-PCs revealed that differentiated hPSC-PCs are most similar to late juvenile (P21) mouse PCs, confirming the efficacy of the protocol in generating more differentiated PCs. Detailed comparison of global gene expression patterns of mouse PCs, using metagene projection analysis, showed that the key gene expression pathways of differentiated hPSC-PCs most closely matched those of late juvenile, mouse PCs (P21). Comparative bioinformatics identified classical PC gene signatures as well as novel mitochondrial and autophagy gene pathways during the differentiation of both mouse and human PCs. In addition, we identified genes expressed in hPSC-PCs but not mouse PCs and confirmed protein expression of a novel human PC marker, CD40LG, expressed in both hPSC-PCs and native human cerebellar tissue. This study therefore provides the first direct comparison of hPSC-PC and mouse PC gene expression and a robust method for generating differentiated hPSC-PCs with human-specific gene expression for modeling developmental and degenerative cerebellar disorders.

A wide range of studies have shown the importance of glia to hPSC-neuronal differentiation (46). For developing mouse PCs, Bergmann glia have been shown to grow in alignment with PC dendritic arbors suggesting a role for glia in PC development (8). Here we observed that co-culturing hPSC-PCs with cerebellar glia prior to culturing them with presynaptic neurons, appeared to promote differentiation as the cells expressed higher levels of the PC marker CALB1 and projected more complex dendrites after co-culture with glia. In addition, cell-cell interactions with their target neurons, GCs, also appeared to promote PC differentiation. As discussed below, functional synaptic connections were necessary for hPSC-PC activity. These findings are consistent with the general observation that co-culture of CNS neurons with glia and with target neurons promotes differentiation.

Expression of the genetically encoded calcium indicator jRGECO1a in hPSC-PCs was an efficient means to measure calcium currents in hPSC-PCs. Our observation that hPSC-PCs co-cultured with mouse GCs fired calcium transients that could be blocked by TTX or CNQX is consistent with the interpretation that the hPSC-PCs were functionally active and had pharmacological properties of PCs. Many of the larger calcium responses showed a slow decay of 30-90 seconds, suggesting release of calcium from intracellular stores (Figure S1C). This classical observation has been made in mouse PCs in cerebellar slices either by simultaneous activation of the climbing fiber and parallel fiber (47) or following train stimulation of parallel fibers only (48) and suggests strong activity in our cultures. In agreement with studies using calcium imaging in *ex vivo* slices, the dynamics of calcium imaging in our co-cultures of hPSC-PCs and mouse GCs did not reveal simple spike activity. However, our results are consistent with the work of others (49) showing the utility of jRGECO1a to image PC somatic calcium, which provides a readout of average PC firing properties.

One of the most consistent observations of hPSC-induced CNS neurons is the relatively slow and often incomplete maturation of human neurons (50). Our results suggest a number of factors likely slow the development of human neurons. On a morphological level, the number of presynaptic inputs varies between human and mouse neurons. This feature is especially dramatic for PCs given the fact that the ratio of GC:PC inputs in human is almost 20 fold higher than mouse (51). This provides one possible explanation for the relatively immature dendritic arborization pattern of the hPSC-PCs we studied. In our experiments, by morphology, hPSC-PCs after ∼17 weeks of differentiation resembled 18 week human PCs (27). It will be interesting to determine whether bio-engineering approaches to increase the density of GC:PC inputs also promote hPSC-PC dendritic density. Another possible reason hPSC-PCs are more immature is the fact that we cultured the cells for four months, while native, human PCs mature over a 30-month period. Finally, the timing of expression of the thyroid hormone gene pathway (*THRA1, RORα*, and *PPARGC1A*), which were shown by Mason and colleagues to rapidly induce mouse PC differentiation beginning at P0 (38) are not expressed until +95 days *in vitro* for hPSC-PCs, suggesting a possible molecular basis for delayed maturation of human PCs.

In contrast to immature morphology we observed for hPSC-PCs, bioinformatic analysis of gene expression in hPSC-PCs and developing mouse PCs, showed that hPSC-PCs most closely resembled late juvenile P21 mouse PCs. This finding suggests that the hPSC-PCs we studied are among the most mature hPSC-derived CNS neurons analyzed to date. This underscores the ultility of transcriptomic analysis for analyzing the maturation of hPSC-derived CNS neurons. Our finding of shared PC signature gene sets in both mouse PC and hPSC-PC transcriptomes validates our conclusion of the identity of hPSC-PCs as Purkinje neurons.

Prior studies using metagene projection analysis have shown the efficacy of this approach for identifying the most salient features of gene expression pathways and for enabling cross-platform and cross-species analysis of gene expression in cancer (36) and in developing CNS neurons, including cerebellar GCs (37). In the present study, metagene analysis of global gene expression patterns of mouse PCs revealed classical PC signatures as well as novel mitochondria and autophagy pathways. These patterns were recapitulated in hPSC-PCs, suggesting that much of human and mouse PC development is conserved.

The finding that hPSC-PCs upregulated mitochondrial gene sets after the formation of synapses with GCs and emergence of calcium firing properties is consistent with a role for mitochondria in calcium buffering and local, activity driven protein synthesis in large output neurons (52). The importance of mitochondria to PCs has been well demonstrated in mouse models and is related directly or indirectly to many autosomal recessive cerebellar ataxias (53). Our findings identify P0-P7 as the time when mitochondrial genes are highly upregulated during mouse PC development and suggest that hPSC-PCs will be useful for studying human PC mitochondrial disorders.

The expression of autophagy gene sets by hPSC-PCs is also important for studies on degenerative cerebellar disorders. PCs are particularly affected in mice conditionally mutant for autophagy genes *Atg5* or *Atg7* (54, 55). In humans, a handful of congenital disorders of autophagy have been described with mutations in *SNX14*, *ATG5*, and *SQSTM1/p62* affecting the cerebellum (56). Like mitochondria genes, our findings define P0-P7 as the time when autophagy genes are upregulated during mouse PC development. The coincidence in timing of mitochondrial and autophagy gene upregulation raises the hypothesis that one key role for autophagy in PCs may be turnover of mitochondria, or mitophagy. Recent studies have found pathological roles for mitophagy in neurons in general and PCs specifically (57, 58). Upregulation of autophagy genes in hPSC-PCs suggests that this model system will be useful for studying human PC autophagy disorders.

Translational profiling of hPSC-PCs and mouse PCs captured species-specific gene expression, consistent with findings of differential gene expression between human and rodent cerebellar cells (59). Two types of species-specific gene expression were observed, expression of primate-specific genes in hPSC-PCs that are non-existent in mouse, and upregulation of genes in hPSC-PCs compared to background levels of expression in mouse PCs. Of the primate-specific genes, the roles of two genes in human neocortex development have recently been studied. *NOTCH2NL* is a human-specific paralog of the NOTCH receptor arising from gene duplication and contributing to expansion of cortical progenitors (39–41). *GLUD2* is a hominoidea-specific, intronless paralog of *Glud1*, arising from retroposition and contributing to changes in cortical gene expression and metabolism (42, 43). The roles of *NOTCH2NL, GLUD2*, and additional primate specific genes in human Purkinje cells is unknown.

Of the genes expressed in hPSC-PCs but not expressed by mouse PCs, we found a sodium channel (*SCN7A*), a regulator of cardiac calcium flux (*PLN*), two genes with known roles in the immune system (*CD40LG* and *GIMAP4*) and a gene of unknown function (*CCDC178*). We further confirmed human-specific gene expression of CD40LG by immunofluorescence. CD40LG is well known for its expression in activated T cells (60) and a potential role in cortical neurite growth has been described (61). Further studies will be necessary to elucidate the function of this novel marker in human PCs and whether it serves a canonical “immune-like” or alternative role.

This study underscores the importance of genetic analysis of mouse and human CNS neurons for modeling CNS disease. Transcriptional profiling is a powerful methodology for confirming the identity of specific CNS neurons as well as for a quantitative measure of their state of differentiation. Future comparative studies on mouse and human neuronal transcriptomics and proteomics will provide insight on improving model systems for CNS neurological disease. To aid in disease modeling, human specific features discovered in our study and others can be introduced into the mouse for more faithful recapitulation of human disease. These “humanized” mice will provide a complimentary system to human pluripotent stem cells.

## Materials and Methods

### CONTACT FOR REAGENT AND RESOURCE SHARING

Further information and requests for resources and reagents should be directed to and will be fulfilled by the Lead Contact, Mary E. Hatten.

### EXPERIMENTAL MODEL AND SUBJECT DETAILS

#### Mice

Wild type C57BL/6J mice (The Jackson Laboratory) were used for isolation of neonatal cerebellar glia and granule cells (P5-P7, see below for details). Isolated cells from both male and female pups were pooled. *Tg(Pcp2-L10a-Egfp)* TRAP mice were a kind gift from N. Heintz (The Rockefeller University, also available from The Jackson Laboratory, B6;FVB-Tg(Pcp2-EGFP/Rpl10a)DR168Htz/J) (32). For isolation of Purkinje cell TRAP RNA, cerebella were pooled from both male and female mice.

All mice were healthy and were housed with a companion(s) in a specific-pathogen free animal facility in vented cages on a 12 hour light/dark cycle. Food and water were provided *ad libitum*. All mouse experiments were performed in accordance with The Rockefeller University IACUC approved protocols.

#### Human ESC Culture and Transgenesis

The RUES2 female human embryonic stem cell line, a kind gift from A. Brivanlou (The Rockefeller University), was maintained in mTESR1 (Stem Cell Technologies, 85850) on hESC-qualified matrigel (Corning, 354277) at 37°C with 5% CO_2_ and sub-cultured weekly with EDTA (Versene, ThermoFisher Scientific, 15040066) (62). Following karyotype analysis, cells were banked in mTESR1 with 10% DMSO in liquid nitrogen. Frozen stocks were thawed into mTESR1 with 10μM Y27632 (Stemgent, 04-0012-02) for the first day. Cells were used within five passages before thawing a new stock to avoid karyotype abnormalities. The RUES2-Pcp2-TRAP cell line was generated using lentiviral transduction (see below for details). All human embryonic stem cell studies were carried out in accordance with the Tri-Sci Embryonic Stem Cell Oversight Committee (ESCRO; Weill-Cornell Medical College, Memorial-Sloan Kettering Cancer Center, The Rockefeller University) approved protocols.

### METHOD DETAILS

#### Generation of RUES2-Pcp2-TRAP hESC Line

Mouse pL7-mGFP (Pcp2-mGFP) plasmid was a kind gift from J. Hammer (National Institutes of Health) (63). pC2-EGFP-L10a was a kind gift from N. Heintz (The Rockefeller University). To subclone the L7 promoter into pC2-EGFP-L10a, oligonucleotides were used to insert a KpnI restriction site into the AseI restriction site of pC2-EGFP-L10a. The KpnI-AgeI fragment of pL7-mGFP containing ∼1kb upstream through part of exon 4 of the mouse Pcp2 gene was subcloned into pC2-EGFP-L10a, replacing the CMV promoter with L7. A PCR product was amplified from pL7-EGFP-L10a using F: 5’-TTCAAAATTTTATCGATTAAGCTTCTCAGAGCATGGTCAG-3’ and R: 5’-AATAGGGCCCTCTAGATTATCTAGATCCGGTGGATCCC-3’ primers containing ClaI and XbaI restriction sites. The PCR product was cloned into the linearized lentiviral vector pSIN-EF1a-promoter-BHH polyA-PGK-Puromycin, digested by ClaI and XbaI, using the In-Fusion HD Cloning Kit (Takara, 638909).

Lentivirus was produced by Lipofectamine 2000 (ThermoFisher, 11668030) transfection of ∼30×10^6^ HEK293FT cells (ThermoFisher, R70007) with psPAX2 (6ug), pMD2.G (2.5ug), and pSIN-L7-EGFP-L10a (7ug). Media was collected and filtered through a 0.45μm PVDF filter after 48-60 hours, concentrated using the Lenti-X concentrator (Clontech, 631231), and stored at −80°C.

Rues2 were dissociated to single cells and seeded at 70,000 cells/well of a Matrigel coated 12 well plate in mTESR1 with 10μM Y27632 (Stemgent, 04-0012-02). The next day 10ul of lentiviral particles were added to fresh mTESR1 medium containing 10μM Y27632 and 4μg/ml protamine sulfate (MilliporeSigma, P3369-10G). After 12 hours the cells were washed with PBS and fed with fresh mTESR1 with 10μM Y27632. After 48 hours, transformed cells were selected by including 1ug/ml puromycin (ThermoFisher, A1113803) in mTESR1 with 10μM Y27632. After an additional 48 hours of selection, cells were dissociated to single cells and 1000 cells were seeded in a 10cm dish in mTESR1 with puromycin and Y27632 and maintained until colonies formed. Clonal lines were manually harvested and checked for normal karyotype (Memorial Sloan Kettering Cytogenetics Core) and for proper EGFP-L10a expression in Pcp2^+^ cells by immunofluorescence following differentiation. RUES2-Pcp2-TRAP lines were maintained in mTESR1 with puromycin when in the undifferentiated state with daily media changes. Puromycin was removed at the start of differentiation.

#### hESC Differentiation

RUES2 maintained in mTESR1 were dissociated into single cells using TrypLE (ThermoFisher, 12605010) and seeded in DMEM/F12 (ThermoFisher, 10565-018) with 1X NEAA (ThermoFisher, 11140050), 1X N2 (ThermoFisher, 17502048), 1X B27 (ThermoFisher, 12587010), 2μg/ml heparin (MilliporeSigma, H3149), 1X Primocin (Invivogen, ant-pm-1), 10mM nicotinamide (MilliporeSigma, N0636), 50ng/ml noggin (Peprotech, 120-10C), 1.5μM CHIR99021 (Stemgent, 04-0004-02), 10 μM Y27631 (Stemgent, 04-0012-02) at 4,000 cells/25μl/well in an untreated 96 well v-bottom plate (ThermoFisher, 12-565-481). Cells were centrifuged for 5 minutes at 1000rpm to promote aggregate formation. For optimal differentiation efficiency, pluripotent cultures should be free of all differentiation, at 50-75% confluence, and cultured for 6-8 days after last passage. It should be noted that mTESR1 contains LiCl, a GSK3β inhibitor, that likely activates some level of Wnt signaling. If undifferentiated cells are cultured in a different pluripotency medium, the concentration of CHIR99021 may need to be altered for proper midbrain/hindbrain specification.

On day 2 of differentiation, 175μl/well of DMEM/F12 with 1X NEAA, 1X N2, 1X B27, 2μg/ml heparin, 1X Primocin, 10mM nicotinamide, 10ng/ml noggin, 1.5μM CHIR99021 was added to the 96 well plate. On day 4 and 5, 175μl/well of medium was replaced with 175μl/well DMEM/F12 with 1X NEAA, 1X N2, 1X B27, 2μg/ml heparin, 1X Primocin, 100ng/ml FGF8b (Peprotech, 100-25).

On day 6, aggregates are plated on laminin coated (ThermoFisher, 27015; 10ng/ml coated overnight at 37 °C) 6-well tissue culture plates or laminin/poly-d-lysine coated glass coverslips (NeuVitro, GG-12-1.5-pdl). Media (DMEM/F12 with 1X NEAA, 1X N2, 1X B27, 2μg/ml heparin, 1X Primocin, 100ng/ml FGF8b) was changed at 2ml/well for 6 well plates or 0.5ml/well 12 well plates with 12mm coverslips. Full media changes were performed on day 8 and 10 using the same media composition and volume.

On day 12, the basal medium was changed to neurobasal (ThermoFisher, 21103049), 1X Glutamax-I (ThermoFisher, 35050061), 1X N2, 1X B27, 1X Primocin, 10ng/ml BDNF (Peprotech, 450-02). Media was subsequently changed every other day using the same composition until isolation of Purkinje cells.

After 22-28 days of differentiation, postmitotic PCs were isolated and co-cultured with mouse cerebellar glial cells. To isolate PCs, differentiated cultures were dissociated by incubating with 0.6mg/ml Papain (Worthington Biochemical, LS003118) in CMF-PBS (dPBS ThermoFisher, 14190-250; 0.2% glucose MilliporeSigma, G8769; 0.004% sodium bicarbonate MilliporeSigma, S8761-100ML; 0.00025% Phenol Red MilliporeSigma, P0290) with 0.23mg/ml L-cysteine (MilliporeSigma, C8277) for 30 minutes at 37°C. Cell clumps were allowed to collect in the bottom of conical tubes and excess papain was removed. 250μl 0.5mg/ml DNAse (Worthington Biochemical, LS002139; in BME ThermoFisher, 21010-046 with 0.33% glucose) was added and clumps were incubated at 37°C for 5 minutes. Cell clumps in DNAse were gently triturated with 3 decreasing bore sizes of fire-polished Pasteur pipettes until mostly single cells. Dissociated cells were passed through a 40μm cell strainer (ThermoFisher, 352340), washed with 10ml BME with 10% horse serum (ThermoFisher, 26050-088), and centrifuged for 5 minutes at 1100rpm.

Cells were resuspended in 5ml CMF-PBS with 3% BSA (MilliporeSigma, A9576-50ml) and plated on anti-GD3 (Biolegend, 917701) 10cm plates (untreated plates coated overnight at 4°C with 13μg GD3 antibody in 10ml 50mM Tris-HCl pH9.5; 1 10cm GD3 plate per 6 well dish of differentiated stem cells; washed 3X PBS before using) and incubated at room temperature for 20 minutes. Plates were tapped to dislodge partially attached cells and the supernatant was transferred to another GD3 plate for an additional 20-minute room temperature incubation. Plates were tapped to dislodge partially attached cells and the supernatant was collected and centrifuged for 5 minutes at 1100rpm.

Cells were resuspended in 5ml neural differentiation medium (BME, ThermoFisher, 21010-046; 1X N2; 1X B27; 0.9%BSA, MilliporeSigma, A9576-50ml; 0.9% glucose, MilliporeSigma, G8769; 008% NaCl; 30nM T3, MilliporeSigma, S8761; 1X Primocin) with 10ng/ml BDNF and incubated at 37°C, 5% CO_2_, for 1 hour. This step allows re-expression of NCAM1, which is cleaved by papain. Cells were collected and centrifuged for 5 minutes at 1100rpm.

Cells were resuspended in 90μl CMF-PBS with 3% BSA and 10μl CD56 (NCAM) microbeads (MiltenyiBiotec, 130-050-401) per 10^7^ cells and incubated at 4 °C for 15 minutes. 10ml CMF-PBS with 3% BSA was added to wash and cells were centrifuged at 1100rpm. Cells were resuspended in 500μl CMF-PBS with 3% BSA and 10μl DNase and applied to a MACS column per manufacturer’s directions (MiltenyiBiotec, 130-042-201). Eluted cells were plated as 90μl droplets at 1×10^5^ cells/cm^2^ on poly-d-lysine coverslips (Neuvitro, GG-12-1.5-pdl; coated overnight with 10ng/ml laminin) with 1×10^4^ mouse cerebellar glia/cm^2^ in 24 well plates. Following attachment (∼1 hour), 1ml neural differentiation media with 10ng/ml BDNF was added per well and coverslips were carefully flipped using fine forceps so that cells were between the bottom of the plate and the coverslip. Mixed cultures were maintained at 35°C with 5% CO_2_. Cultures were 0.5ml media was exchanged once per week. After 65-70 days, coverslips were gently flipped right side up and 1×10^6^ mouse cerebellar granule cells/cm^2^ were added. 0.5ml media was subsequently exchanged every Monday and Friday, with 4μM Ara-C (MilliporeSigma, C6645-100MG) added 1 week after granule cell addition.

#### Isolation of mouse cerebellar glia and granule cells

Mouse cerebellar glia and granule cells were isolated as previously described (64). Briefly, cerebella from P5-P7 mice were dissected, dissociated to single cells with trypsin and Dnase (Worthington Biochemical 3703 and 2139), and separated on a 35%/60% percoll gradient resulting in two layers of cells. The top layer contained glia, interneurons, and PCs. This layer was collected and plated on Matrigel in DMEM/F12 (ThermoFisher, 10565-018) with 1X NEAA (ThermoFisher, 11140050), 1X N2 (ThermoFisher, 17502048), 1X B27 (ThermoFisher, 12587010), 2μg/ml heparin (MilliporeSigma, H3149), 1X Primocin (Invivogen, ant-pm-1) with 10% horse serum (ThermoFisher, 26050-088). Glia were grown and passaged at least once before use to remove neurons and were cultured for up to two passages before co-culture with hPSC-PCs as described above. The bottom layer contained granule cells and where added to hPSC-PC/glia cultures as described above.

#### Quantitative Real-Time PCR

RNA was isolated using the RNeasy Plus mini kit (Qiagen, 74134). cDNA was generated from 1μg RNA using iScript cDNA Synthesis Kit (Bio-Rad, 1708891). qPCR was carried out on a Roche LightCycler 480 (Roche, 05015278001) using the SYBR green method (Roche, 04707516001) in triplicate 10μl reactions run in a 96-well plate using half the cDNA synthesis reaction per plate. The qPCR protocol was: 95°C 5 minutes; 45 rounds of: 95°C 15 seconds, 56°C 15 seconds, 72°C 10 seconds; followed by a melt curve analysis from 65°C-95°C. Primer specificity was confirmed by melting temperature analysis and gel electrophoresis. Data were normalized to the geometric mean of the “housekeeping” genes: glyceraldehyde phosphate dehydrogenase (*GAPDH*), hydroxymethylbilane synthase (*HMBS*), and glucose phosphate isomerase (*GPI*) (65). Primer sequences are listed in Supplementary table 1.

#### Immunofluorescence Labeling

Cells were fixed with 4% paraformaldehyde (Electron Microscopy Sciences, 15710) in dPBS (ThermoFisher, 14190-250) for 10 minutes at room temperature. In some instances, samples were embedded in O.C.T. medium (Electron Microscopy Sciences, 62550-01) and sectioned on a Leica CM3050S cryostat. Mouse cerebella and thymus were fixed in 4% paraformaldehyde in dPBS overnight at 4°C and cut into 50μm sections on a Leica VT1000S vibratome.

Samples were blocked in dPBS with 5% BSA and 1% donkey serum (Jackson Immunoresearch, 017-000-001). Concentration of primary antibodies and % Triton X-100 are listed in the Supplementary table 2. Primary antibodies were incubated overnight at 4°C. After washing with PBS, appropriate secondary Alexa Fluor antibodies (Thermofisher) were incubated at 1:500 for 1 hour at 4°C. In some instances, DAPI was added for 5 minutes at room temperature before final washing in PBS. Coverslips were mounted in Fluoro-Gel (Electron Microscopy Sciences, 17985-10) and sealed with nail polish. Images were captured on an inverted Zeiss Axio Observer Z1 laser scanning confocal microscope.

7μm paraffin sections of human 5-day cerebellum and human tonsil were rehydrated and treated with Trilogy solution (Cell Marque, 920P-04) in a conventional steamer for 40 minutes, followed by 20 minutes cool down at room temperature. Sections were blocked for 2 h at room temperature in 10% normal goat serum, 1% BSA, 0.1% TritonX-100 in PBS, pH 7.4 and then incubated overnight at 4°C with mouse monoclonal anti-CD154 (1:50, R&D systems, MAB617) and rabbit anti-calbindinD28k (1:500, Swant) in antibody diluent (1% BSA, 0.1% TritonX-100 in PBS, pH 7.4). Sections were washed x3 in PBS-0.1% Tween, followed by incubation for 2 hours at room temperature in antibody diluent with anti-mouse and anti-rabbit goat secondary antibodies conjugated to Alexa Fluor 568 and Alexa Fluor 488, respectively (Invitrogen, Carlsbad, CA). After three washes in PBS-0.1% Tween, sections were mounted in DAPI Fluoromount-G (SouthernBiotech, 0100-01). Samples were collected at Columbia University Irvine Medical Center, New York, NY with previous patient consent and in strict accordance with institutional and legal ethical guidelines.

#### TRAP-RNA Isolation

TRAP-RNA was isolated as previously described (30, 37, 66). Briefly, for *Tg(Pcp2-L10a-Egfp)* mice, pooled cerebella were immediately homogenized with a Teflon-glass homogenizer in ice-cold polysome extraction buffer (10 mM HEPES-KOH [pH 7.4], 150 mM KCl, 5 mM MgCl2, 0.5 mM DTT (MilliporeSigma, D9779-1G), 100 μg/ml cycloheximide (MilliporeSigma, C7698-1G), Superasin and RNasin RNase inhibitors (ThermoFisher, AM2694, PR-N2515)and complete-EDTAfree protease inhibitor (MilliporeSigma, 11836170001)). Following clearing by centrifugation, supernatants were incubated at 4°C with end-over-end rotation for 16-18 hours with biotinylated Streptavidin T1 Dynabeads (ThermoFisher, 65601) previously conjugated with GFP antibodies (Sloan Kettering Institute Antibody Core, HtzGFP-19C8 and HtzGFP-19F7). The beads were collected on a magnetic rack, washed, and resuspended in lysis buffer with β-mercaptoethanol (Agilent, 400753) to extract bound RNA from polysomes. RNA was purified using the Stratagene Absolutely RNA Nanoprep kit (Agilent, 400753). For RUES2-Pcp2-TRAP hPSC-PCs, polysomes were stabilized by adding 100 μg/ml cycloheximide to cell culture media for 10 minutes prior to homogenization with polysome extraction buffer and isolation as described above. RNA was purified using the RNeasy micro kit (Qiagen, 74004). RNA quantity and quality was measured using an Agilent 2100 Bioanalyzer with the 6000 Pico Kit (Agilent, 5067-1513).

#### Microarray

Microarray experiments were performed as previously described (37). Briefly, purified RNA was amplified using the SuperScript GeneChip Expression 3’-Amplification Reagents Two-Cycle cDNA Synthesis Kit (Affymetrix, Santa Clara, CA) and the GeneChip T7-Oligo Primer (Affymetrix, Santa Clara, CA). The cDNA was used for the in vitro synthesis of cRNA using the MEGAscriptT7 Kit (Ambion, Austin, TX). cRNA was purified using the GeneChip Sample Cleanup Module (Affymetrix, Santa Clara, CA). 600 ng or less of clean cRNA was 29 used in the second-cycle cDNA synthesis reaction using the SuperScript GeneChip Expression 3’-Amplification Reagents Two-Cycle cDNA Synthesis Kit (Affymetrix, Santa Clara, CA) and random primers (Affymetrix, Santa Clara, CA). The cDNA was purified using the GeneChip Sample Cleanup Module (Affymetrix, Santa Clara, CA). Purified cDNA was used for the in vitro synthesis of biotin labeled cRNA using the GeneChip IVT Labeling Kit (Affymetrix, Santa Clara, CA). cRNA was purified using the GeneChip Sample Cleanup Module (Affymetrix, Santa Clara, CA) and fragmented into 35-200 base pair fragments using a magnesium acetate buffer (Affymetrix, Santa Clara, CA). Ten micrograms of labeled cRNA were hybridized to Affymetrix GeneChip Mouse Gene 1.0 ST Array for 16 h at 45°C. The GeneChips were washed and stained according to the manufacturer’s recommendations (Affymetrix, Santa Clara, CA) using the GeneChips Fluidics Station 450 (Affymetrix, Santa Clara, CA). Mouse Gene 1.0 ST arrays were scanned using the GeneChip Scanner 3000 (Affymetrix, Santa Clara, CA).

#### RNA Sequencing

1 ng of total RNA was used to generate full length cDNA using Clontech’s SMART-Seq v4 Ultra Low Input RNA Kit (Cat # 634888). 1 ng of cDNA was then used to prepare libraries using Illumina Nextera XT DNA sample preparation kit (Cat # FC-131-1024). Libraries with unique barcodes were pooled at equal molar ratios and sequenced on Illumina NextSeq 500 sequencer to generate 75 bp single reads, following manufactures protocol (Cat# 15048776 Rev.E)

#### Calcium Imaging

To image calcium spikes in hPSC-PCs, post-mitotic cells were nucleofected during the isolation step of the differentiation protocol (after 22-28 days). Following GD3 negative immunopanning, the semi-purified mixture of cells was nucleofected with 2μg/10^6^ cells of each: pL7-mGFP ((Pcp2-mGFP) kind gift from J. Hammer (National Institutes of Health) (63)) and jRGECO1a (pAAV.Syn.NES-jRGECO1a.WPRE.SV40, kind gift from D. Kim & GENIE Project, Addgene plasmid #100854, (29)). Nucleofection was performed on an Amaxa nucleofector IIb using protocol O-003 with the mouse neuron kit (Lonza, VPG-1001). Cells were allowed to recover for 1 hour at 37 °C and followed by NCAM1 MACS and culture as described above.

Following isolation and nucleofection, cells were cultured with mouse cerebellar glia for 70 days and then mouse cerebellar granule cells for an additional 30 days. To record changes in fluorescence over time, cells expressing both jRGECO1a and GFP were recorded by an Andor iXon 512×512 EMC camera using an inverted Zeiss Axiovert 200 with a Perkin-Elmer UltraView spinning disk confocal head. Images were captured at 3.47hz for a total of 143.712 seconds per recording. Baseline recordings on Day +100 were made from 23 cells from 3 separate culture dishes. 25μM CNQX was added and recordings were made from 15 cells from 3 separate culture dishes. CNQX was washed out and media was replaced. On day 101 baseline recordings were made to confirm activity before adding 300nM tetrodotoxin (Tocris, 1078). Recordings were made from 10 cells from 3 separate culture dishes. Tetrodotoxin was washed out and resumption of calcium activity was confirmed.

To analyze recordings, files were opened with ImageJ (NIH) and integrated density from an individual cell was analyzed over time. Fo was set as the lowest integrated density over the recording. Traces of ΔF/Fo in Figure 1I and Supplemental Figure 1C are representative traces of various calcium firing patterns we observed. Traces in Figure 1J/K are representative traces in the presence of synaptic inhibitors. No spikes were ever observed in the presence of synaptic inhibitors.

### QUANTIFICATION AND STATISTICAL ANALYSIS

#### qPCR Analysis

Three biological replicates for each differentiation condition were analyzed. Relative gene expression levels were normalized to the geometric mean of the “housekeeping” genes: glyceraldehyde phosphate dehydrogenase (*GAPDH*), hydroxymethylbilane synthase (*HMBS*), and glucose phosphate isomerase (*GPI*) (65). Student’s T-test was used for statistical comparison between two groups using GraphPad Prism software.

#### Microarray Analysis

Microarray data was normalized using RMA methods implemented in Affymetrix Power Tools (67) and changes between functional groups assessed using the Limma Bioconductor package (68). Clustering of mouse transcriptome profiles was performed using Non-negative Matrix Factorization implemented in the NMF package (69). Single sample GSEA analysis was performed using the GSVA Bioconductor package (70) with query gene-sets retrieved from MsigDB databases (71) or derived as GMT format from the Human RNA-seq data.

#### RNA Sequencing Analysis

Sequence and transcript coordinates for human hg19 UCSC genome and gene models were retrieved from the Bioconductor Bsgenome.Hsapiens.UCSC.hg19 and TxDb.Hsapiens.UCSC.hg19.knownGene libraries respectively. Unaligned sequence reads were retrieved as FastQ format. FastQ files for Day24 samples were downsampled using the ShortRead R package to equilibrate total mapped reads to Day95 samples. Transcript expression was calculated using the Salmon software quantification (72) and gene expression levels as TPMs and counts retrieved using Tximport (73). Differential gene expression analysis was performed using DESeq2 (74). For visualisation in genome browsers, RNA-seq reads are aligned to the genome using Rsubread’s subjunc method (75) and exported as bigWigs normalised to reads per million using the rtracklayer package.

To further assess any potential contamination by mouse feeder cells following human specific IP, RNAseq data was aligned to an In-Silico Combined Genome (ISCG) and a conservative, mouse subtracted, human transcriptome dataset was acquired (34). To create the ISCG, sequence and transcript coordinates for mouse mm10 UCSC genome and gene models were retrieved from the Bioconductor Bsgenome.Mmusculus.UCSC.mm10 and TxDb.Mmusculus.UCSC.mm10.knownGene libraries and combined with the above hg19 genome. RNAseq reads were then mapped to the ISCG with Subread and gene expression estimates acquired using Featurecounts and Salmon. Human specific gene expression estimates were then retrieved from the ISCG estimates and used in differential gene expression performed with DEseq2.

## ACKNOWLEDGEMENTS

We are grateful to members of the Hatten lab for helpful discussions; the Brivanlou lab and Stem Cell core (The Rockefeller University) for helpful discussions and reagents; the Heintz lab (The Rockefeller University) for helpful discussions and reagents; Dr. John Hammer (NIH) for reagents; Dr. Pablo Tamayo (University of California, San Diego) for helpful discussion; Dr. Mustafa Sahin (Boston Children’s Hospital) for helpful discussion; summer students Maya Pandit, Lauren Cantor, and Victoria Marino; and The Rockefeller University Bio-Imaging, Genomics, Comparative Bioscience Center, and Bioinformatics resource centers. This work was Supported in part by grant #UL1 TR001866 from the National Center for Advancing Translational Sciences (NCATS, National Institutes of Health (NIH) Clinical and Translational Science Award (CTSA) program) administered by The Rockefeller University Center for Clinical and Translational Science; by NIH/NINDS grant 1R21NS093540-01; by DOD grant W81XWH-15-1-0189; and by a Starr Tri-Institutional Stem Cell Initiative grant from the Starr Foundation.

