## Supplemental Figures 1-4 and Supplemental Tables 1-2 for "Novel genetic features of human and mouse Purkinje cell differentiation defined by comparative transcriptomics"

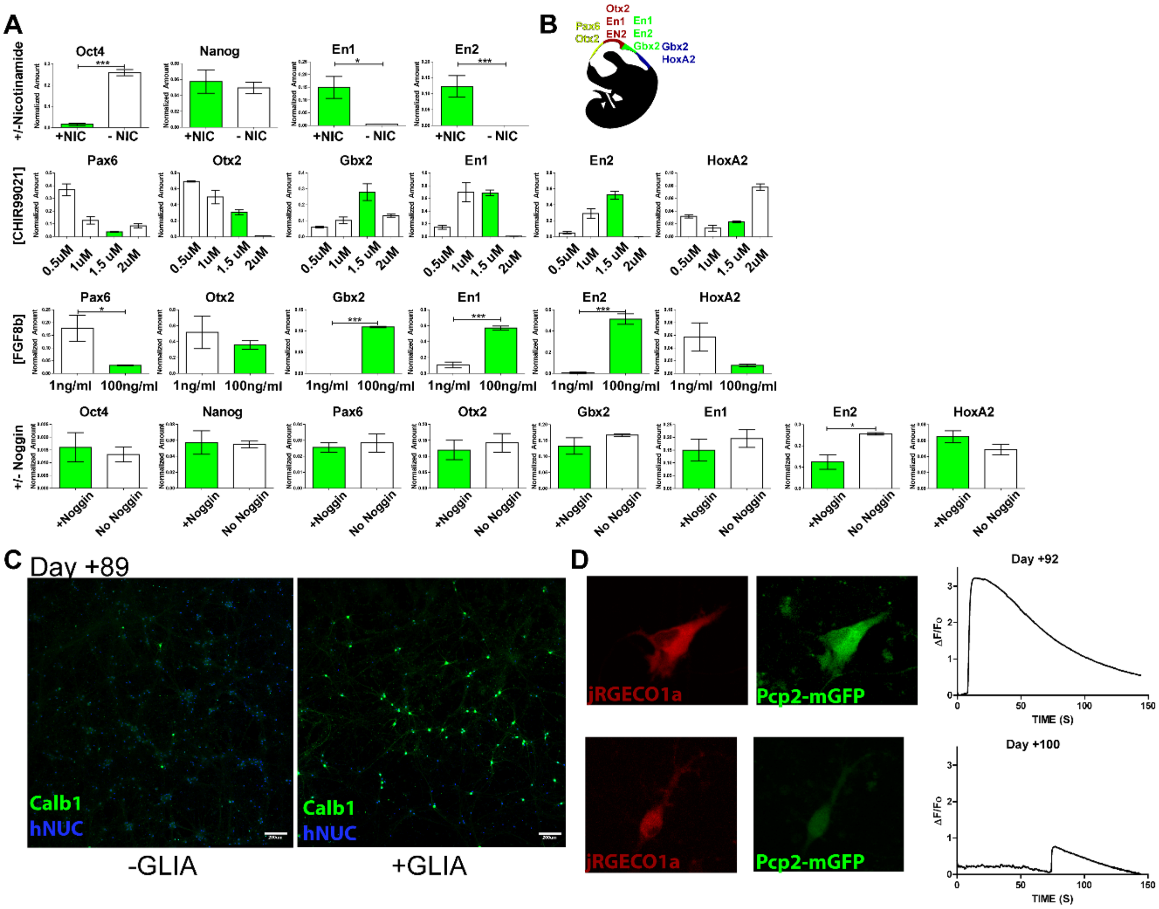


**Supplemental Figure 1. Differentiation of hPSCs to PCs. A.** qPCR analysis at day 6 of differentiation in the presence or absence of differentiation factors. Green bars represent optimized conditions used subsequently. Student's T-Test P-values *<0.05 ***<0.001. Data were collected from three rounds of differentiation and error bars show standard error of the mean. **B.** Schematic of neurodevelopmental patterning depicting rostral-caudal marker genes. Cerebellar genes and the cerebellar primordium are in green. **C.** Live imaging of genetically encoded calcium indicator jRGECO1a and hPSC-PC reporter Pcp2-mGFP after 92 and 100 days of co-culture with mouse glia and granule cells and traces of the change in fluorescence (ΔF/Fo) over time.

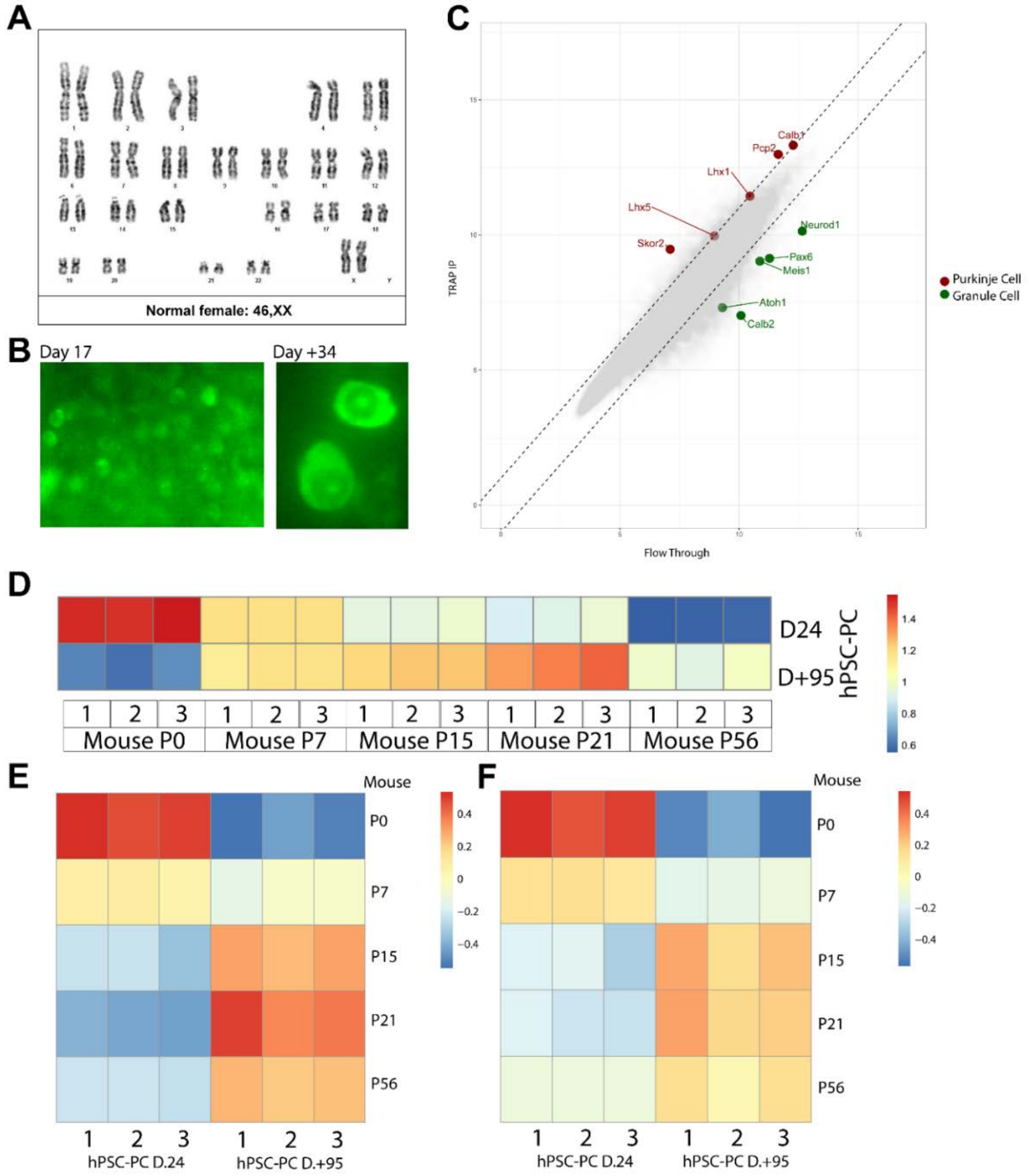


**Supplemental Figure 2. Translational Profiling in hPSC-PCs and Mouse PCs. A.** Karyotype analysis of RUES2-PCP2-L10a-EGFP stem cell line. **B.** Live imaging of the L10a-EGFP TRAP reporter after 17 days of differentiation and an additional +34 days of differentiation following isolation and co-culture. **C.** Test for enrichment of PC markers and depletion of GC markers from TRAP isolation of RNA from a P7 *Tg(Pcp2-L10a-Egfp)* TRAP mouse, dotted lines represent $\text{Log}_2$ fold change. **D.** Heat map depicting expression levels in mouse PCs over postnatal development of gene sets defined as $\log_2$ four-fold change between Day 24 and Day +95 in hPSC-PCs following subtraction of mouse genes from hPSC-PC data using an in silico human-mouse reference genome (34). Day 24 hPSC-PCs are most similar to P0 mouse PCs ($p=5.26\times10^{-17}$). Day +95 PCs are most similar to P21 mouse PCs ($p=3.73\times10^{-5}$). **E.** Heat map depicting expression levels in hPSC-PCs of mouse gene sets defined as the 100 most expressed genes per time point. **F.** Heat map depicting expression level in hPSC-PCs of mouse gene sets defined as the 100 most expressed genes per time point following subtraction of mouse genes from hPSC-PC data using an in silico human-mouse reference genome (34).

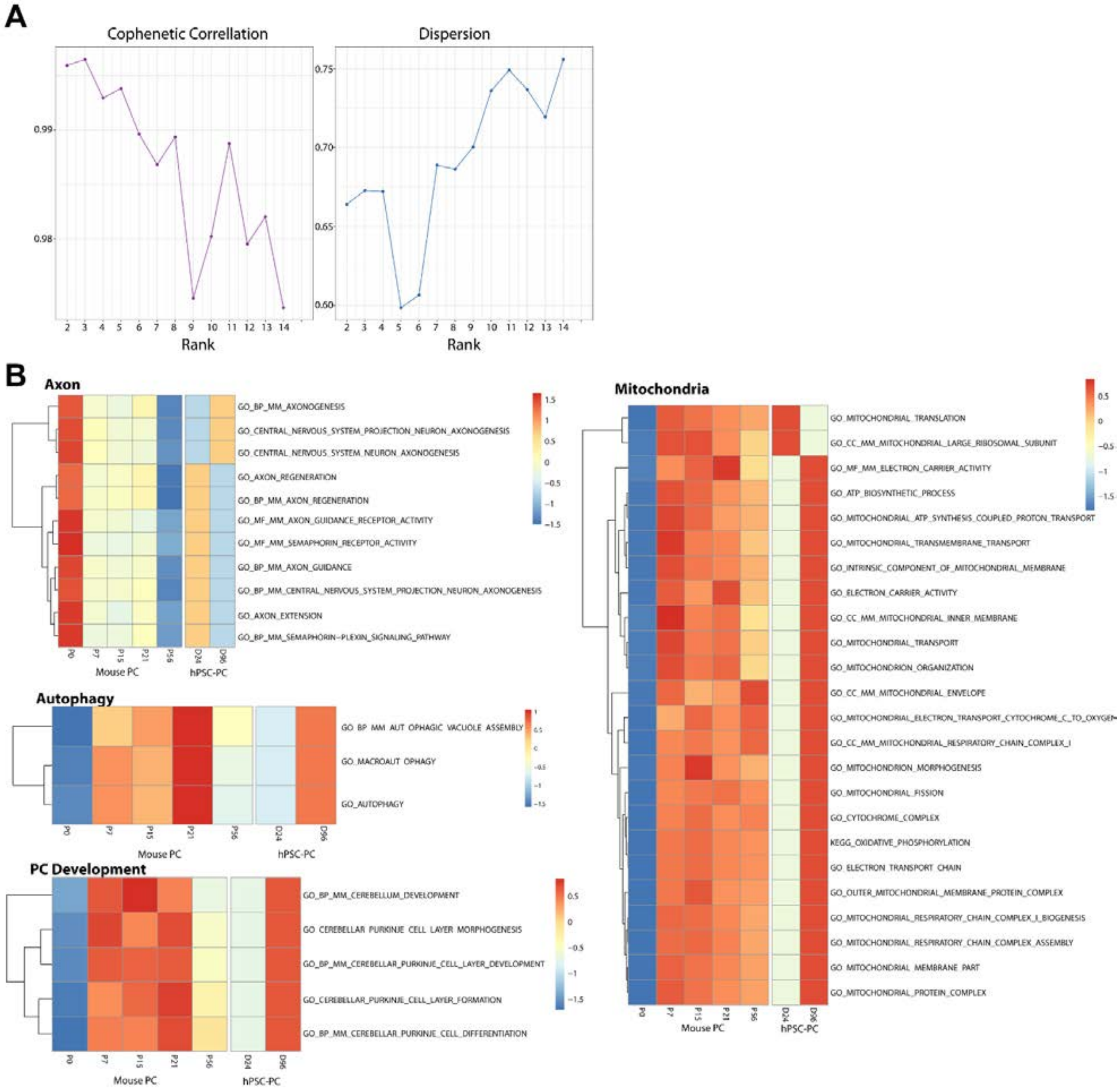


**Supplemental Figure 3. NMF metagene analysis of mouse PC TRAP Data. A.** Cophenetic correlation and dispersion by rank (metagene number). Rank 5 showed high cophenetic correlation with low dispersion. **B.** Heat maps depicting all gene ontology terms for the gene ontology signatures (Axon, Autophagy, PC Development, Mitochondria) in Figure 3.

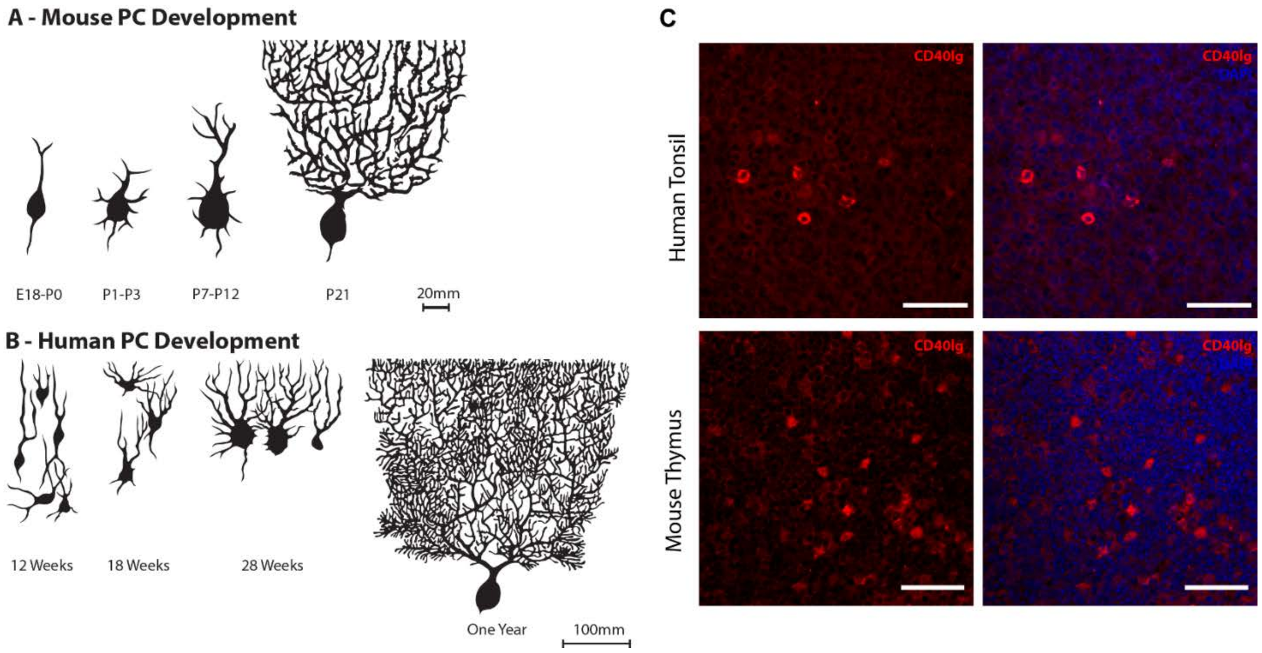


**Supplemental Figure 4. Species differences. A.** Schematic of mouse PC development. (Adapted from (76)and (77)). **B.** Schematic of human PC development (Adapted from(27)). **C.** Positive controls for anti-human CD40LG antibody and anti-mouse CD40LG antibody. Scale bars = 50μm.

| Gene | Forward | Reverse | Source |
|---|---|---|---|
| Oct4 | CAGTGCCCGAAACCCACAC | GGAGACCCAGCAGCCTCAAA | (78) |
| Nanog | CAGAAGGCCTCAGCACCTAC | ATTGTTCCAGGTCTGGTTGC | (78) |
| Pax6 | TCACCATGGCAAATAACCTG | CAGCATGCAGGAGTATGAGG | Designed on Primer3 |
| Otx2 | ACAAGTGGCCAATTCACTCC | GAGGTGGACAAGGGATCTGA | Designed on Primer3 |
| Gbx2 | GTTCCCGCCGTCGCTGATGAT | GCCGGTGTAGACGAAATGGCCG | Designed on Primer3 |
| En1 | GAGCGCAGGGCACCAAATA | CGAGTCAGTTTTGACCACGG | Primerbank (126090908c1) |
| En2 | GGCGTGGGTCTACTGTACG | TACCTGTTGGTCTGGAACTCG | Designed on Primer3 |
| HoxA2 | CGTCGCTCGCTGAGTGCCTG | TGTCGAGTGTGAAAGCGTCGAGG | (24) |
| GPI3’ | GGACCACGAGCCCTTAGC | AACACTTCAGCCAATTCTAACAC | (65) |
| GPI5’ | CGTCATCAACATTGGCATTGG | GGGACCTCCTGAAGAGTATGG | (65) |
| HMBS | TGCTATCTGGGGAGTGATTACC | GGCTGTTGCTTGGACTTCTC | (65) |
| GAPD | AGCAAGAGCACAAGAGGAAGAG | GAGCACAGGGTACTTTATTGATGG | (65) |

**Supplementary Table 1. qPCR Primer Sequences.**

| Antibody | % Triton X-100 | Dilution | Source |
|---|---|---|---|
| En1 | 0.3 | 1:10,000 | Gift of T. Jessell |
| Otx2 | 0.3 | 1:2000 | Millipore Cat# AB9566, RRID:AB_2157186 |
| Kirrel2 | 0.3 | 1:250 | R&D Systems |
| Corl2(Skor2) | 0.3 | 1:200 | Atlas Antibodies Cat# HPA046206, RRID:AB_2679588 |
| Pcp2(L7) | 0-0.1 | 1:500 | Takara Cat#M202 |
| Calb1 | 0.3 | 1:250 | Swant Cat# 300, RRID:AB_10000347 |
| Calb1 | 0.3 | 1:500 | Swant Cat# CB38, RRID:AB_2721225 |
| GD3 | 0 | 1:200 | BioLegend Cat# 917701, RRID:AB_2565200 |
| NCAM(CD56) | 0 | 1:200 | BioLegend Cat# 304601, RRID:AB_314443 |
| hNUC | 0.3 | 1:200 | Millipore Cat# MAB1281, RRID:AB_94090 |
| Syn1 | 0.3 | 1:500 | Sigma-Aldrich Cat# S193, RRID:AB_261457 |
| GFP | 0.3 | 1:4000 | Aves Labs Cat# GFP-1020, RRID:AB_10000240 |
| Human CD40lg | 0.05 cell culture, 0.1 paraffin | 1:50 | R and D Systems Cat# MAB617, RRID:AB_2291414 |
| Mouse CD40lg | 0.05 | 1:50 | R and D Systems Cat# AF1163, RRID:AB_35463 |

**Supplemental Table 2. Antibody List.**
